# Gene editing in the Chagas disease vector *Rhodnius prolixus* by Cas9-mediated ReMOT Control

**DOI:** 10.1101/2023.08.14.553172

**Authors:** Leonardo Lima, Mateus Berni, Jamile Mota, Daniel Bressan, Alison Julio, Robson Cavalcante, Vanessa Macias, Zhiqian Li, Jason L. Rasgon, Ethan Bier, Helena Araujo

## Abstract

*Rhodnius prolixus* is currently the model vector of choice for studying Chagas disease transmission, a debilitating disease caused by *Trypanosoma cruzi* parasites. Despite the broad use of double-stranded RNA interference for the knockdown of gene function in *R. prolixus*, transgenesis and gene editing protocols are still lacking. Here we tested Receptor-Mediated Ovary Transduction of Cargo (ReMOT Control) and direct parental injection of CRISPR (DIPA-CRISPR) for the maternal delivery of CRISPR/Cas9 elements to developing *R. prolixus* oocytes, and strategies for the identification of insertions/deletions (indels) in target loci of resulting gene-edited G0 nymphs. We demonstrate successful ReMOT Control-mediated gene editing of the eye color markers *Rp-scarlet* and *Rp-white*, and the cuticle color marker *Rp-yellow,* with highest effectiveness obtained using the ovary-targeting BtKV ligand. These results provide proof-of-concepts for generating somatic mutations in *R. prolixus* and potentially for generating germline edited lines in triatomines. Our studies also suggest that optimal strategies for recovery of mutations include performing multiple gRNA injections and the use of visible phenotypes such as those displayed in the *Rp-scarlet, Rp-white* and *Rp-yellow* loci for future Co-CRISPR experiments. These results will lay the foundation for gene editing protocols for triatomines and could lead to the development of novel control strategies for vectors of Chagas disease.

**Author Summary:** *Rhodnius prolixus* is an insect vector of the protozoan *Trypanossoma cruzi*, causative agent of debilitating Chagas disease. To fight the spread of the disease, it has been suggested that biological control of the insect should be attempted. Gene editing by the novel CRISPR methodology holds great promise in this sense, as it enables to target almost any gene in the genome for mutagenesis, thus allowing the control of insect physiology and reproduction. Here we have tested protocols for the delivery of CRISPR reagents as an attempt to enable genome editing of the vector. Our results show that maternal delivery of CRISPR by the ReMOT Control method is efficient for mutating eye and cuticle color genes in the resulting nymphs, generating edited animals with red eyes, white eyes or a yellow cuticle. This is the first report of gene editing in a vector of Chagas disease and should lay the basis to produce modified animals either unable to carry the *T. cruzi* parasite or to reproduce.

## Introduction

Chagas disease is a neglected tropical disease that inflicts great health and socioeconomic impacts in the Americas. Caused by the protozoan parasite *Trypanosoma cruzi*, reduviid insects represent one of the most common transmission routes, either by the insects delivering parasites during blood feeding or by oral ingestion of the parasite-carrying insects with açai fruit [1]. Chagas disease treatments are limited, however, and vector control is essential to restrict spread of the disease. The kissing bug *Rhodnius prolixus* is the most well studied laboratory species among the triatomines that transmit *T. cruzi*. With a sequenced genome [2], gene identification and transcriptome analyses have revealed important aspects of *R. prolixus* physiology and evolution [3–6]. Gene knockdown by parental RNA interference (pRNAi) is effective in *R. prolixus*, allowing investigation of gene function [7–9]. Such studies would be greatly enhanced, however, by advances in transgenic and gene-editing technologies, which could enable efficient functional genetic studies and potentially lead to the development of novel biological control strategies for the insect vectors.

CRISPR/Cas9 (clustered regularly spaced palindromic repeats/Cas-9 associated) methods have revolutionized precise genome editing in a broad range of animal lineages. These effective and widely used tools benefit from its simplicity and the great number of putative target sequences for practically every gene. Adapted from bacteria, the most common technique involves complexing the Cas9 endonuclease with single guide RNAs (sgRNAs), which recognize target sequences in the genome and direct site-specific double stranded breaks (DSBs) by the action of the Cas9 nuclease [10]. These DSBs are then corrected by a series of repair mechanisms, such as Non-Homologous End-Joining (NHEJ) or Homology Directed Repair (HDR) [11]. NHEJ is error-prone and often generates short insertions and deletions (indels) at the cut site, leading to loss-of-function mutations. HDR, on the other hand, enables the precise insertion of exogenous sequences at the cut site, permitting facile site-directed modification of target sequences and insertion of transgenic gene cassettes.

Most animal CRISPR protocols rely on the injection of sgRNA/Cas9 complexes or plasmids containing sequences for the production of CRISPR elements into the embryo by embryonic microinjection. These injections are performed during early embryonic development when nuclei are easily accessible, increasing the chance of gene edition in the germline. Experiments of this type are quite challenging to perform in triatomines such as *R. prolixus*. First, oogenesis in these animals is strictly dependent on blood ingestion [12]. 3-5 days after blood feeding, vitellogenic oocytes start to develop and females lay approximately 30-40 eggs in the next three weeks. Therefore, collecting embryos at the appropriate stage for injections (0-2h development) requires a large number of ovipositing animals. Second, eggs are encapsulated by a hard chorion [13] that is extremely difficult to penetrate for injection with DNA constructs, resulting in low viability of injected embryos. Alternative methodologies that bypass the need for embryonic microinjections have been developed, which involve injecting CRISPR components into the adult female hemolymph followed by uptake of these components by developing oocytes [14–18]. While such methods do not yet allow the incorporation of exogenous sequences into target sites by HDR, they have been shown to generate small deletions and base alterations by NHEJ that may produce effective gene knockouts.

Previously, we identified visible markers that would serve as appropriate target sites for gene editing in *R. prolixus*. Through RNA interference we identified several loci that when disrupted lead to visible cuticle or eye color phenotypes with little or no effect on insect fertility or viability. Such viable and fertile phenotypes are essential for generating edited lines and following the heritability of editing events through multiple generations [19]. Here we tested two techniques: Receptor-Mediated Ovary transduction of Cargo (ReMOT Control, [14]) and Direct Parental CRISPR (DIPA CRISPR, [17]) for delivery of Cas9/gRNA complexes to oocytes and generation of gene edited offspring. We show that while DIPA-CRISPR was not effective in *R. prolixus*, ReMOT Control was able to generate CRISPR-edited G0 offspring for several targeted cuticle and eye color genes. These results are the first to accomplish gene editing in triatomine bugs, opening possibilities for the development of germline gene editing strategies to address functional genetic questions in triatomines, and applied strategies for Chagas disease control.

## Methods

### Insect rearing

*R. prolixus* rearing was performed at 28°C and 70-75% humidity, with animals fed on rabbit blood throughout all developmental stages. Animal care and experimental protocols were conducted following guidelines of the Committee for Evaluation of Animal Use for Research from the Federal University of Rio de Janeiro (CEUA-UFRJ, #01200.001568/2013-87, order number 155/13). Technicians dedicated to the animal facility at the Institute of Medical Biochemistry (UFRJ) conducted all aspects related to rabbit husbandry under strict guidelines to ensure careful and consistent animal handling.

### Injections to determine ovary intake of P2C- and BtKV-tagged fluorescent proteins

To determine whether P2C sequences and BtKV sequences were efficient to deliver proteins to *R. prolixus* oocytes, we injected young adult females with untagged mCherry, P2C-mCherry and BtKV-mCherry into the abdomen (1μl, 2,5 or 5 μg/μl). As control for background fluorescence we injected females with the same buffer used for mCherry proteins. Tagged and untagged mCherry protein was produced as in [14] (provided by J. Rasgon). Subsequently, animals were blood fed and 2-3 days later the ovaries were dissected, fixed in paraformaldehyde 4%, mounted in 70% glycerol and observed under a Leica MZ10F Fluorescent Stereomicroscope. To test whether P2C and BtKV were efficient for oocyte delivery in other Chagas disease vectors, we obtained *Triatoma infestans* animals from the International Reference laboratory for Triatomine Taxonomy at Fundação Oswaldo Cruz, Rio de Janeiro, injected the females and analyzed the oocytes as above. To determine the amount of fluorescence uptake, we used the Analyze function in Fiji, collecting the mean grey value for a constant area in vitellogenic oocytes, since mature chorionated oocytes display a great amount of background fluorescence.

### sgRNA design and synthesis

sgRNAs were designed against *Rp-y*, *Rp-w*, *Rp-sca* and *Rp-aaNAT* loci in order to target sequences at the beginning of the coding region or close to enzyme activation sites using CHOP CHOP [20]. sgRNAs were either synthesized at Synthego or synthesized in loco with Invitrogen Megascript^tm^ T7 Transcription (AM1334) [21], using the primers in Table S1 and S2.

### gRNA/Cas9 injections

gRNA/Cas9 complexes were prepared as in [14]. Briefly, *R. prolixus* females were fasted and injected with a solution containing 5 μg Cas9 and 2.5 μg of gRNA. Cas9 proteins used were P2C-Cas9 [14] and BtKV-Cas9 [16], prepared as above by J. Rasgon lab, or commercially purchased Cas9 (PNABIO). The injection was performed between the third thoracic and the first abdominal segment, using a 10ul Hamilton microsyringe adjusted so as not to exceed 2μl of solution for each adult female. The microsyringe was cleaned before and after each injection with hot water, 70% alcohol and finally milliQ water, to avoid possible contamination of the animals during the procedure. After the injection, the insects were kept at rest for one day and then fed using rabbit blood. Females were observed for 15 days to obtain information on survival and egg laying. Injections were performed prior blood feeding, since those performed after blood feeding resulted in a great number of female deaths.

### *in vitro* cleavage assays

To ensure the effectiveness of designed gRNAS, we performed *in vitro* cleavage assays (Figure S2) with gRNA/Cas9 complexes as in [21]. Briefly, the ability of each sgRNA to direct Cas9 cleavage of the target DNA was tested *in vitro* by checking the integrity of the target DNA region after incubation of: PCR amplified and purified target sequences (with Monarch^®^ PCR & DNA Cleanup Kit T1030), sgRNA (30 ng; 3.2 μM) and Cas9 protein (400 ng; 3.8 μM). After incubation and Cas9 inactivation the DNA was analyzed by electrophoresis to search for the presence of fragments resulting from cleavage of the double-stranded DNA.

### Quantification of fecundity and embryonic viability

Eggs were collected from 2 to 3 weeks after blood ingestion, counted, and the hatch rate defined after 20 days at 28°C. To define possible effects of gRNA/Cas9 injections on fecundity, in some experiments females were placed in individual vials for egg collection. Embryonic viability was defined by counting the number of hatched embryos per total number of eggs laid.

### Phenotypic screening

Injected females were reared individually (particularly for multiple sgRNA injections) or in groups of 4-5 females. All first instar nymphs (>48h after hatching) were screened for cuticle or eye color phenotypes using a dissecting scope, under a stream of CO_2_. Animals were fed and kept to develop throughout all stages in an attempt to cross the animals displaying a visible phenotype. However, none of the animals displaying visible phenotypes survived the six months required to reach adulthood. A subset of G1 brothers and sisters of the animals presenting visible phenotypes were sequenced to check for putative edition events in heterozygosity.

### Image acquisition

Images displaying visible phenotypes were obtained using a Leica MZ10F Stereomicroscope, with animals sleeping under a stream of CO_2_.

### Genomic DNA extraction and PCR Protocol

Genomic DNA was extracted from G0 nymphs using a modification of the single-fly method developed for *Drosophila melanogaster* [22]. Briefly, DNA was extracted from single first nymphs or nymph exuvia by macerating in a 50μl solution containing 49μl of “squishing buffer” (10mM Tris-Cl pH 8.0; 1mM EDTA; 25mM NaCl) and 1μl of proteinase K (20mg/ml). After maceration in solution, an incubation period is carried out at 37°C for 20 minutes for enzymatic action, followed by five minutes at 95°C for enzyme inactivation. At the end of the process, the liquid part was transferred to a new tube in order to discard the exoskeleton debris. The DNA-containing solution was kept at -20°C. Target sequences were PCR amplified using primers flanking the gRNA target site (Table S3) and either sent for Sanger or for Illumina sequencing as detailed below. For animals presenting visible phenotypes PCR products were also cloned into TOPO TA cloning vector (Invitrogen) for transforming NEB 5-alpha Competent Cells (E. coli C2987P). At least 10 colonies per condition were analyzed. Plasmid DNA was isolated from each colony under standard protocols and purified with QIAprep Spin Miniprep Kit (Cat. No. / ID: 27104) for Sanger sequencing.

### Sanger Sequencing

PCR products or plasmid DNA were used for Sanger sequencing at the Genomic DNA sequencing facility at the Institute of Biophysics Carlos Chagas Filho at UFRJ. Sequencing results were aligned to reference sequences using Snapgene. At least 10 wild type animals were used to sequence around each gRNA target site to enable the detection of putative SNPs.

### Amplicon Deep Sequencing

To identify and quantify mutations at the target loci, sequences around the gRNA cleavage site were amplified by PCR from G0 or G1 nymph genomic DNA using Q5 Hot-Start High Fidelity Polymerase. PCR products were purified, and 20μg/μl of each sample was sent for Illumina paired-end Amplicon-based deep sequencing at Genewiz. All reads were analyzed using CRISPResso2 (https://crispresso.pinellolab.partners.org/submission) with wildtype sequences as reference. Primer sequences and amplicon sizes for each target are detailed in Table S3.

### Statistical analyses

Statistical analyses were performed through unpaired t test with Welch’s correction. Differences were considered significant at *p*<.05. All statistical analyses were performed using the Prism 7.0 software (GraphPad Software).

## Results

### ReMOT delivery of proteins to Triatomine ooctyes

We tested whether ReMOT Control could deliver components to triatome oocytes *in vivo* by first injecting 4-7 day old *R. prolixus* females with mCherry protein fused to either the P2C- or the BtKV-oocyte delivery sequences [14,16], and measured the amount of fluorescence in developing oocytes. Unmodified mCherry was injected as a control. The P2C peptide is derived from the N-terminal portion of *D. melanogaster* Yolk Protein 1 and has been shown to increase the delivery of sgRNA/Cas9 complexes to oocytes in multiple Dipteran, Hymenopteran, Coleopteran, and Hemipteran species [15,23,24]. The BtKV peptide is derived from the *Bemisia tabaci* vitellogenin binding sequence and has been shown to deliver cargo to vitellogenic oocytes in the hemipteran *Bemisia tabaci* [16].

Following injections into *R. prolixus* oocytes, we observed that unmodified mCherry could be taken up to some extent when injected into the hemolymph, but that oocyte uptake levels were significantly increased when mCherry was fused to either the P2C or BtKV ovary targeting sequences. Furthermore, we found that the BtKV peptide was more effective than P2C based on quantitation of fluorescence intensity in the ovaries (Figure 1). To test the generality of these observations, we also injected untagged and tagged forms of mCherry into female *Triatoma infestans* (another important vector of Chagas disease). We observed similar results as those in *R. prolixus* (Figure 1F-H), indicating that P2C- and BtKV increase oocyte delivery of molecular cargo in multiple triatomine vectors.

**Figure 1.**
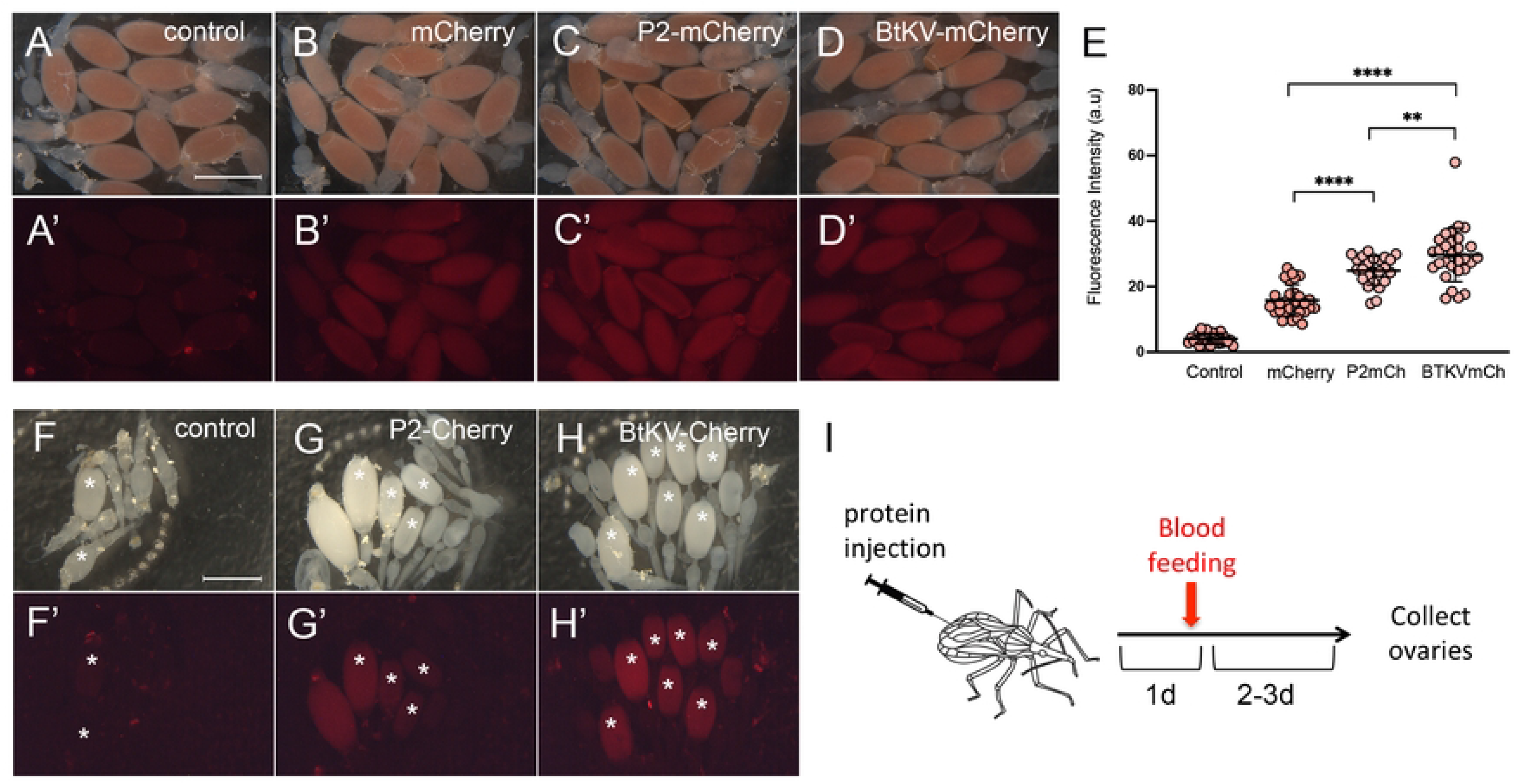
ReMOT tagged proteins are efficiently transferred to triatomine vitellogenic oocytes. A-D) Dissected ovaries from *Rhodnius prolixus* adult control females (A), or females injected with mCherry (B) P2C-tagged mCherry (C) or BtKV-tagged mCherry (D) protein, 3 days after blood feeding. Light microscopy (A-D) and fluorescence (A’-D’) show that fluorescence is greater in tagged proteins. E) Quantification of fluorescence in vitellogenic oocytes. Statistical analysis performed through unpaired t test with Welch’s correction. \*\**p*<0.005, \*\*\*\**p*<0.0001. F-H) A similar pattern is observed in *Triatoma infestans* dissected ovaries from adult control females (F), or females injected with P2C-tagged mCherry (G) or BtKV-tagged mCherry (H) protein seen through light (F-H) or fluorescence microscopy (F’-H’). Asterisks indicate vitellogenic oocytes. I) Injection protocol for ovary collection and analysis. Scale bars: 2mm.

### CRISPR/Cas9 editing for genes encoding visible markers in *R. prolixus*

Next, we tested conditions for gene editing in *R. prolixus*, using maternal delivery of sgRNA/Cas9 RNP complexes. We designed sgRNAs targeting coding sequences of genes that encode cuticle or eye pigmentation proteins, expecting that loss-of-function (lof) mutations in these genes should produce visible phenotypes [19]. The targeted loci were: *Rp-yellow* (*Rp*-*y*), which encodes an enzyme required for the synthesis of melanin during cuticle pigmentation; *Rp-scarlet* (*Rp-sca*), which encodes an ABC transporter that transports pigments into the eye; *Rp-white* (*Rp-w*), which encodes a second ABC transporter that transports pigments into the eye; and *Rp-aaNAT*, which encodes an enzyme required for cuticle pigmentation (Table S1). sgRNAs designed for the different loci were injected as preformed ribonucleoprotein complexes with either unmodified Cas9 (DIPA-CRISPR) [17], or with P2C-Cas9 [14] or BtKV-Cas9 [16] to test ReMOT Control. Most injections included the endosomal escape reagent chloroquine, as it has been shown to increase ReMOT effectiveness in some systems (Figure S1). Injections performed before blood feeding have no significant effect on survival or on the number of eggs laid per female [19].

Among all injections performed, we were able to identify animals displaying visible phenotypes in G0 nymphs for complexes targeting the *Rp-y*, *Rp-sca* and *Rp-w* loci, but observed no obvious phenotypes for the *Rp-aaNAT* locus (Table 1). In contrast, no visible phenotypes were observed for any of these loci in nymphs following DIPA-CRISPR injections. Among the animals injected with the ReMOT Control method, we observed several offspring displaying red eyes (instead of the wild-type black eye phenotype) when mothers were injected with *Rp-sca* gRNA, one individual displaying white eyes derived from a mother injected with *Rp-w* gRNA, and one individual (*y29*) displaying a yellow cuticle obtained from a mother injected with *Rp-y* gRNA. For *Rp-sca*, the eye phenotype was seen either by injecting a single sgRNA for *Rp-sca* or when co-injected with other gRNAs. Total DNA prepared from a subset of G0 nymphs that displayed no visible phenotypes was subjected to PCR for amplification of the regions surrounding the sgRNA target site followed by Sanger sequencing. No gene editing events were detected by this method. Nymphs with visible phenotypes were followed each day in an attempt to maintain the animals for genetic crosses as adults. However, these animals were among the nymphs that did not survive to the next stage and DNA was extracted from these individuals upon death and prepared for DNA sequence analysis.

**Table 1.** CRISPR editing efficiency in G0.

| Cas9 protein | gRNA | females injected (n) | G0 nymphs screened (n) | visible phenotype (% , n) | G0 nymphs sequenced*** (n) | G0 with altered sequence (n)**** |
| --- | --- | --- | --- | --- | --- | --- |
| P2C-Cas9 | yellow 1 | 10 | 116* | 0 | nd | - |
| P2C-Cas9 | scarlet 1 | 10 | 85* | (nd; 1) | nd | - |
| BtKV-Cas9 | yellow 1 | 4 | 148 | yellow cuticle<br>(0.68%, 1) | 50 | 1 (y29) |
| BtKV-Cas9 | scarlet 1 | 4 | 135 | red eye<br>patches<br>(0.75%, 1) | 33 | 0 |
| BtKV-Cas9 | white 1 | 5 | 142 | white eyes<br>(0.70%, 1) | 20 | 1 |
| BtKV-Cas9 | yellow 2 | 5 | 95 | 0 | 36 | 0 |
| BtKV-Cas9 | aaNAT 1 | 5 | 141 | 0 | 15 | 0 |
| BtKV-Cas9 | scarlet 2 | 5 | 133 | 0 | 45 | 0 |
| BtKV-Cas9 | 4 gRNAs** | 2 | 80 | red eyes<br>(3,95%, 3) | 10 | 1 |
| Cas9 (DIPA) | 4 gRNAs** | 7 | 236 | 0 | nd | nd |
| buffer | NA | 6 | 91* | 0 | nd | nd |

P2C-Cas9 injections were performed in the absence of chloroquine. All other injections were performed with 0.5mM chloroquine.
\*eggs counted for one week only (second week after blood feeding), total nymphs screened exceeded 300 for these conditions.
\*\* A mixture of 4gRNAs were injected with Cas-9 protein: white 1, yellow 2, aaNAT 1, scarlet 2.
\*\*\* Sequencing was performed by Sanger using total genomic DNA
\*\*\*\* Sequencing was performed by Illumina Deep Sequencing and/or Sanger Sequencing using amplified sequences cloned into bacteria

### Confirmation of gene editing events in *Rp-yellow, Rp-white* and *Rp-sca*

Total genomic DNA from one G0 third-instar nymph displaying an entirely yellow cuticle (Figure 2) was extracted and sequences surrounding the *Rp-y* gRNA target site were PCR amplified and cloned into a TOPO TA cloning vector (Invitrogen) for bacterial transformation. Single colonies were Sanger sequenced with M13 or specific primers from both sides of the target site. We detected a large deletion in 3/10 colonies sequenced. This lesion is predicted to generate a truncated and non-functional protein (Figure 2D,E). Importantly, we observed an 11bp deletion located 24bp 3’ of the Cas9 cleavage site in 3/8 control animals and 13/30 injected animals, characterizing heterozygotes for a deletion that is present in the colony. This 11bp deletion resides inside an intron, is not predicted to alter protein function and does not result in a visible phenotype, since animals homozygous for the deletion display no phenotype. Despite apparently not altering gene function, the deletion in heterozygotes precludes the use of heteroduplex assays for the screening of nearby lesions generated by CRISPR.

**Figure 2.**
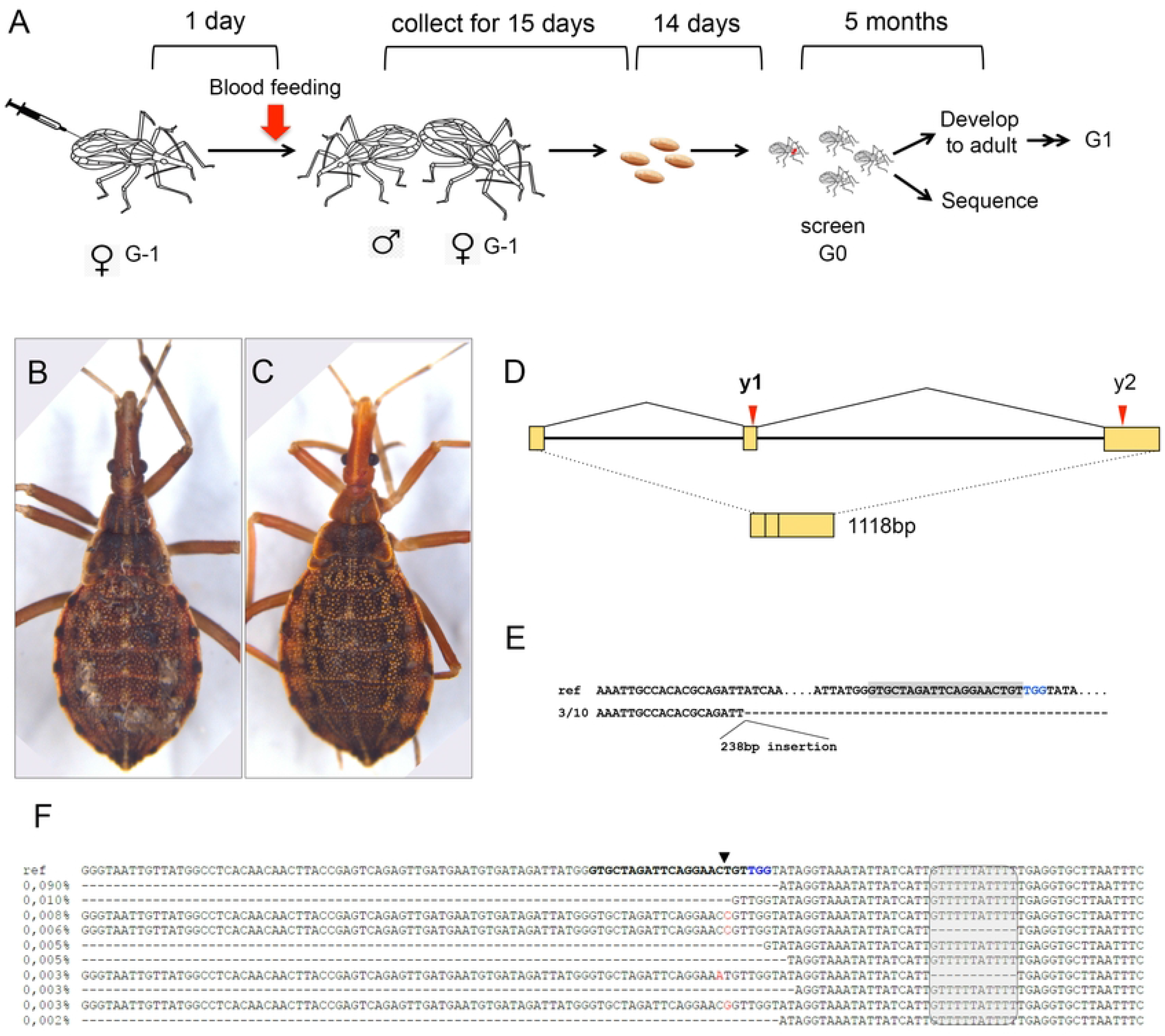
CRISPR-directed gene edition events in the *yellow* locus. A) Protocol for gRNA/BtKV-Cas9 RNP injection into the adult *R. prolixus* female hemolymph. G-1 injected females are mated and eggs are collected in individual vials, allowing define the number of G0 mosaic visible phenotypes in resulting nymphs per injected female. Half G0 animals are let develop and crossed between siblings. The remaining half is used for DNA isolation and amplicon analysis. B,C) Third instar siblings resulting from an RNP injected female showing a wild type (B) and a yellow cuticle (C) phenotype. D) *Rp-yellow* genomic region with exons (boxes) and introns, as well as target region for the two *y* gRNAs tested. In bold the gRNA that resulted in the depicted phenotypes. E) Sequence amplified from the animal in C, cloned into bacteria. In gray the gRNA sequence, in blue the PAM, in the reference sequence. The cloned sequence from C shows a 238bp insertion and a large deletion that covers the gRNA target site. The deletion (4657bp) covers the entire exon 2 (186bp) and part of the intron between exons 2 and 3. F) Amplicon sequencing for the same individual shows with low frequency (percentages) a series of sequences that display deletions or base changes at the target site. The grey box indicates a natural deletion present in heterozygosity. In bold the gRNA sequence, in blue the PAM. Red corresponds to nucleotide substitutions.

The large deletion surrounding the Cas9 cleavage site in the yellow nymph (*y29*) is surprising, since NHEJ events expected to correct DSBs frequently generate small indels. However, large deletions resulting from CRISPR/Cas9 gene editing have been reported in ticks [18]. To determine the different types of lesions resulting from the CRISPR editing process, we amplified and performed next generation deep sequencing of the region surrounding the gRNA target site. Small indels were observed at low frequencies, as well as a series of larger deletions (Figure 2F; Table S4). These deletions are consistent with that observed by Sanger sequencing and indicate that *y29* is a mosaic containing a mixture of wild type *Rp-yellow* alleles and alleles that harbor deletions and base changes surrounding the target site, that likely impair *Rp-yellow* function. Analysis of the sequences surrounding the *Rp-white* gRNA target site in the white-eyed nymph, on the other hand, displayed no large sequence modifications (Figure 3A-D). Sanger sequencing of the nymph genomic DNA revealed no modification relative to controls, but a small indel was detected by PCR amplification and cloning in 1/10 colonies. However, the C>A base change is located 71bp 3’ of the predicted Cas9 cleavage site and may represent a rare SNP. Deep sequencing revealed a small fraction of sequences displaying single nucleotide modifications at the Cas9 cleavage site (Figure 3D; Table S5). The most frequent of these sequences was predicted to encode an amino acid substitution, replacing a tryptophan with a cysteine residue, a significant alteration that could potentially impair protein function (Figure 3E).

**Figure 3.**
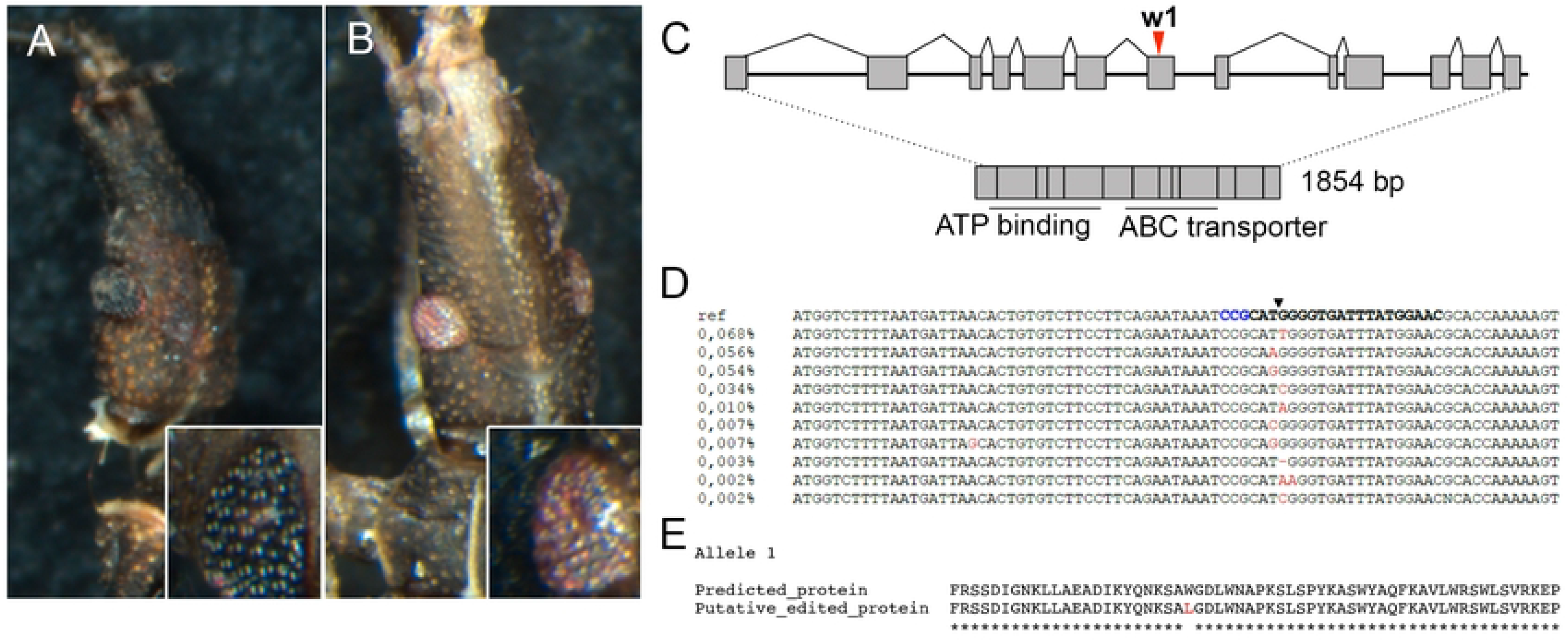
Gene edition in the white locus leads to loss of eye color. A,B) First instar siblings resulting from an RNP injected female showing a wild type (A) and a white eye (B) phenotype. C) *white* locus with gRNA target site. D) Amplicon sequencing for the first instar animal in B shows a series of low frequency indels. E) The most frequent of the base substitutions generates an amino acid replacement that could impair protein function.

Subsequently, we examined nymphs for phenotypes likely resulting from impairment of *Rp-sca* function. Four nymphs displayed a red eye phenotype, one with only slightly red eyes (from P2C-Cas9 injection, and were not analyzed further) and three individuals with strong red eye phenotypes (from independent BtKV-Cas9 injected mothers). One of these three individuals was derived from a single *Rp-sca* gRNA injection and the other two from injections with gRNAs against multiple genes (Figure 4A-C). We chose one of the multiple gRNA-injected nymphs for Illumina deep sequencing. Analysis of the sequences surrounding the *Rp-sca* gRNA target site revealed a significant amount (3.3% of aligned reads) of base substitutions located three to four nucleotides from the PAM site, indicating likely imprecise repair through the NHEJ pathway (Figure 4E). The most frequently observed nucleotide substitution is predicted to result in an amino acid replacement, from an alanine to a negatively charged glutamic acid residue (Figure 4F). Since the gRNA target site is placed in the region that encodes the ATP binding site, this substitution could impair protein function leading to the impaired transport of dark pigments in the eye [19]. Although none of the red-eyed G0 nymphs survived to adulthood, one G1 animal resulting from crosses among the brothers and sisters of the red-eyed G0 (in Figure 4A,C) displayed a mosaic eye color phenotype, wherein one eye was red while other eye had ommatidia as black as the wild type (Figure 4G,H). We did not detect edited alleles by Sanger sequencing, suggesting that the animal might be a mosaic with a small number of edited alleles, arguing against the hypothesis that it resulted from gene editing events transmitted through the germline.

**Figure 4.**
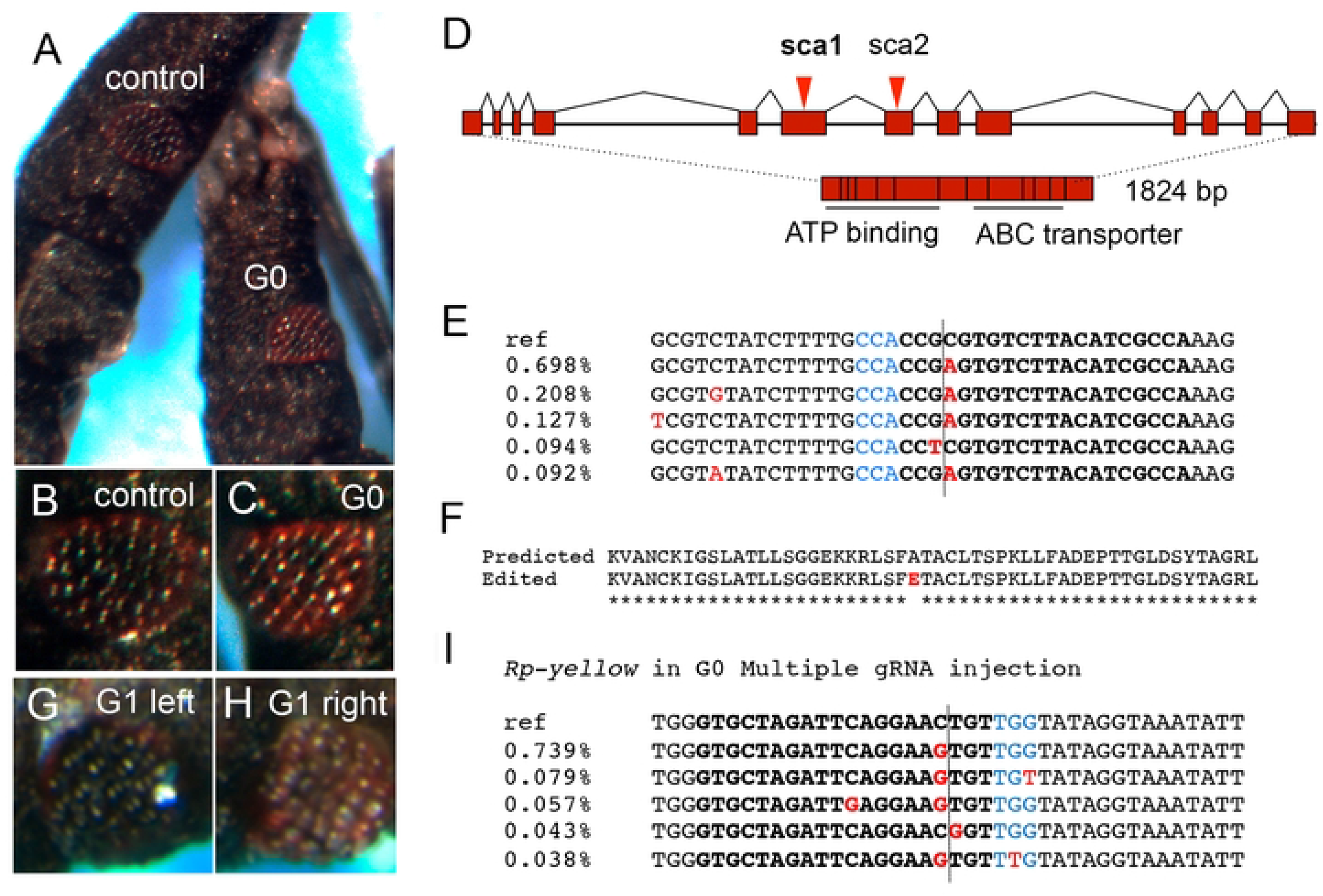
Red eyes result from *scarlet* gRNA/BtKV-Cas9 single and multiple gRNA injections. First instar siblings resulting from a female injected with multiple gRNAs (directed to *Rp-sca, Rp-y, Rp-w and Rp-aaNAT*) in BtKV-Cas9 RNPs, showing a wild type (A,B) and a scarlet eye (A,C) phenotype. D) *Rp-scarlet* locus with gRNA target sites. In bold the gRNA used for the animal depicted in C. E) Amplicon sequencing for the first instar red eyed animal in C. Shown are the most frequent sequences, which display base substitutions positioned 3 nucleotides from the PAM site. F) The frequent C>A base substitution results in an aminoacid residue substitution located inside the ATP binding site of the ABC transporter encoded by *Rp-scarlet*. G,H) A G1 nymph resulting from the cross between brothers and sisters of the G0 shown in C. The animal displays one black (G) and one red (H) eye. I) Amplicon sequencing for the *Rp-yellow* locus in the first instar red eyed animal in C. Several base substitutions in the Rp-y target region are observed. Red in E,I indicate base substitutions, blue indicates the PAM site and the gRNA target sequence is in bold. The dashed line represents the expected PAM cleavage site. The percentage of the representative sequences refers to those defined around the gRNA site only.

Next, we investigated whether a visible phenotype resulting from genome editing in one locus (*Rp-sca*) could help identify co-editing events in other loci that would not generate any visible effect. Therefore, we sequenced the *Rp-w* and *Rp-y* gRNA target sites in the red-eye nymph resulting from the multiple gRNA ReMOT. Base substitutions were detected along a guanine stretch in the *Rp-w* target site, three nucleotides from the PAM, both in the G0 and the G1 nymph. However, a similar percentage of modified sequences were observed in a control nymph that was sequenced under identical conditions, suggesting that these variant alleles did not result from Cas9-mediated editing events (Figure S3). On the other hand, sequence analysis for *Rp-y* in G0 identified several altered sequences (1.84% of reads, as compared to the control 0.41% modified sequences) at the predicted Cas9 cleavage site (Figure 4I; Figure S3), despite the lack of a detectable cuticle phenotype expected to result from a putative loss of *Rp-yellow* function. In addition, G1 *Rp-y* sequencing identified a series of deletions surrounding the gRNA target site (2.58%) that were represented in yet a much smaller percentage in control wild-type sequences (0.06%; Figure S3). We note that the multiple gRNA-injected nymphs analyzed were first instars that were not allowed to develop to further stages. Thus, we could not confirm whether a visible cuticle phenotype resulting from these alterations in the *Rp-y* locus would appear at later stages, similar to what we observed with the y29 third instar nymph shown in Figure 2.

## Discussion

### ReMOT Control is more effective than DIPA-CRISPR in delivering CRISPR cargo to triatomine oocytes

CRISPR/Cas9 mutagenesis has recently emerged as an efficient method for genome engineering in a number of arthropod species. Most methodologies involving CRISPR rely on the embryonic delivery of CRISPR elements by microinjection. However, in animals that lay eggs encased in a hard chorion such as triatomines, or other types of structures that preclude embryonic delivery of CRISPR elements such as the cockroach ootheca, maternal delivery may be a more accessible method for gene edition. Here we tested oocyte delivery of RNP CRISPR complexes containing untagged and tagged Cas9 protein injected into the female hemocoel. First we showed that untagged mCherry protein was able to enter vitellogenic oocytes, but that fusing mCherry to ovary targeting sequences delivered significantly more protein.

Using the DIPA-CRISPR method (untagged Cas9), we did not detect any animals displaying cuticle or eye color phenotypes resulting from injections with untagged Cas9 (7 females injected, 236 nymphs screened), while with the ReMOT Control (using Cas9 fused to two different ovary targeting peptides) we detected multiple editing events in offspring. Optimization of either method may increase the efficiency of genome editing but, currently, our results indicate that ReMOT control is a more effective method than DIPA-CRISPR for genome editing in *R. prolixus*. The greater amount of oocyte fluorescence resulting from hemocoel injection of BtKV-mCherry relative to untagged mCherry in *T. infestans* suggests that this greater efficiency may hold true for other triatomine vectors of Chagas disease.

### Visible markers for genome edition in *R. prolixus*

Pigmentation mutants have been classically used as genetic markers in recombination experiments and genetic crosses since the early days of *Drosophila* genetics [25]. Pigmentation genes are also frequently carried as visible markers in transgenes, such as a forshortened *mini-white* version of the *w* gene in *Drosophila* [26] and kynurenine monooxygenase gene in *Aedes* [27]. Cuticle and eye color loci may also be very useful in co-CRISPR experiments to generate alleles in loci that may not engender visible phenotypes on their own [28,29]. In such cases individuals exhibiting pigmentation phenotypes may have increased chances of displaying lesions in a second locus that is targeted in parallel. Here we have shown that visible phenotypes can be generated by CRISPR-mediated genome editing of pigmentation loci in *R. prolixus*, corroborating previous evidence from gene knockdown studies [19]. Three loci (*Rp-y*, *Rp-w* and *Rp-sca*) display easily scored phenotypes in the eye and cuticle resulting from CRISPR lesions. Visible phenotypes are seen since the first instar for *Rp-w* and *Rp-sca* but may be harder to detect in young *Rp-y* mosaics. For instance, the light color of the y29 animal was only perceived at third larval instar stage, possibly due to maternal provision of *Rp-y* messages obscuring the mutant phenotype in younger first instars. Notwithstanding, the use of a cuticle marker may be best suited to accompany some types of loss-of-function experiments, such as behavioral experiments that require visible cues. Among the four loci tested, no phenotype for *Rp-aaNAT* was observed, despite the striking dark phenotype resulting from knockdown experiments with RNA interference [19]. This may be due to inefficiency of the gRNA, lethality resulting from gene knockout, or to other limitations.

As mentioned above, ReMOT Control delivery of multiple gRNAs indicates that the parallel delivery of marker gRNAs such as that for *Rp-sca* may help identify edition events in loci that are not expected to generate a visible phenotype. This procedure, termed co-CRISPR, has been shown effective in *C. elegans*, based on a worm movement phenotype [30], and in *Drosophila*, where *ebony* [31] and *ovoD* [28] enhanced the screening and identification of genome-editing events. Among the loci herein tested, *Rp-sca* was most effective in generating animals with a visible phenotype. Importantly, *Rp-sca* knockdown does not alter viability [19], indicating that knockouts may be viable as well. Conversely, *Rp-w* KD decreases viability in *R. prolixus* and *Of-w* loss-of-function is lethal in the Hemiptera *Oncopeltus faciatus* [32]. These differences in viability may be irrelevant while screening G0 mosaics but could lead to loss of edited G1 individuals if *Rp-w* were used.

### Gene editing efficiency in the kissing bug parallels that reported for other insects

Two important characteristics define the suitability of gene editing methods for mutagenesis: the efficiency, defined by the number of edited animals in a population, and germline transmission of induced mutations. Due to the long generation time for *R. prolixus* (approximately 6 months) and the limited number of gene edited individuals capable of surviving this protracted period, determining germline transmission is extremely challenging. Therefore, we defined editing efficiency at generation zero (G0), taking into account that animals displaying visible phenotypes are likely mosaics carrying biallelic lesions at target loci.

Considering the different target loci, we find an editing efficiency (number of edited animals in the population) of 0.7-4.0% in the G0s, resulting from ReMOT control with BtKV-Cas9. This is within the range reported for ReMOT delivery in *Aedes aegypti* (1-2.5%; [14]), *Anopheles stephensi* (∼2%, [15]), *Ixodes scapularis* (1.7-4.2%, [18]), *Culex pipiens* (∼2%, [33]) and *Bemisia tabaci* (0.2-2%; [16]), as scored in the G0. It is important to take into account that the editing efficiencies reported for the species above were defined solely based on visible phenotypes. Conforming to this practice, screening total genomic nymph DNA by Sanger sequencing has proven unsuitable to identify gene editing events in *R. polixus*, as it revealed no detectable sequence modifications, even in mosaic animals displaying visible phenotypes. This indicates that the proportion of edited cells in animals displaying visible phenotypes is likely to be small in relation to wild type cells of the entire animal, consistent with the modest percentage of edited sequences revealed by Deep sequencing. Since similar G0 editing efficiencies reported are correlated with germline transmission of gene editing events in several species listed above, future experiments with large numbers of injected *R. prolixus* females may enable recovery of heritable germline transmitted edited alleles in this species.

Due to the long *R. prolixus* life cycle and the limited number of edited animals obtained, we were yet not able to identify germline transmission of edited events. The unexpected identification of a G1 nymph with one sole red eye is suggestive of some type of partial loss-of-function allele transmitted through the germline. However, Sanger sequencing for the entire nymph or for PCR-amplified sequences cloned into bacterial vectors revealed no detectable sequence modifications, indicating that the visible phenotype results from a small number of somatic editing events. If correct, this could imply that Cas9/gRNA complexes can be transmitted from one generation to another, from the BtKV-Cas9 injected mothers (G-1), through G0, to G1. Although surprising, particularly considering the long *R. prolixus* generation time, a similar form of “shadow drive”, wherein Cas9/gRNA complexes are transmitted to progeny in the absence of gRNA or Cas9 transgenes, has been reported in *Drosophila* [29]. Furthermore, we have observed transmission of an expression DNA plasmid for 2 generations post-ReMOT injection in *An. stephensi* (Macias and Rasgon, unpublished). As such enduring effects would have strong implications for biological vector control, future studies should be undertaken to investigate the possibility of transgenerational delivery of gRNA/Cas9 complexes in triatomines.

### Challenges for genome edition in *Rhodnius prolixus*

In addition to their long generation time, we found other challenges for gene editing in *R. prolixus*. One quite unexpected observation was sequence heterogeneity. Despite the long inbreeding of our *R. prolixus* colony, we found multiple deletions carried as heterozygotes in target loci. As discussed above, this characteristic impairs the use of heteroduplex analysis to screen for editing events. Unfortunately, isogenic lines are too weak for use in these types of experiments, indicating that one should sequence many insects before designing gRNAs for gene editing and that sequencing should not be restricted to the gRNA target region, but should also cover any nearby region that will be PCR amplified during screening.

Another challenge will be to define conditions for repair by HDR, in order to enable the precise modification of target sequences and to incorporate exogenous DNA. Currently, ReMOT control only enables gene edition and repair by NHEJ. This limitation could be solved in the future through the development of effective embryo delivery of CRISPR elements or alternative methods including maternal delivery of RNPs associated with exogenous sequences containing homology arms to sequences flanking the Cas9 cleavage site.

## Acknowledgements

We would like to thank Dr. Pedro L. Oliveira for helpful comments on the manuscript. We are grateful to the animal facility at the Institute of Medical Biochemistry for technical assistance with *R. prolixus* husbandry, and to the International Reference Laboratory for Triatomine taxonomy for providing *Triatoma infestans* animals.

## Supporting Information Captions

**Figure S1. Effect of cloroquine injection on egg viability**. We explored putative toxic effects of maternal injections of the endosomal escape reagent cloroquine on embryo viability. Young females were injected in the abdomen with 1μl of cloroquine at the concentrations displayed on the X axis. Eggs were collected and viability defined 20 days after egg lay. Numbers inside bars correspond to the number of eggs collected per condition. Since we observed a decrease in viability with 1.0mM cloroquine, we chose to use 0.5mM for CRISPR assays.

**Figure S2.** *in vitro* cleavage assay for selected gRNAs. We tested the effectiveness of the designed gRNAs by an *in vitro* cleavage assay. PCR fragments surrounding each of the target gRNA sequences were produced and purified for use in an enzymatic reaction carrying the target double stranded PCR fragment, with (+RNP) our without (ctrl) ribonucleoprotein complexes of purified Cas9 plus the corresponding gRNA. Shown are the tests performed for the gRNAs sca2, aaNAT, y1 and w1. All complexes lead to cleavage of the target DNA. Asterisks denote fragments resulting from Cas9 cleavage of the double stranded DNA.

**Figure S3.** Most represented sequences from Amplicon sequencing surrounding gRNA target sites for *Rp-white* and *Rp-yellow* in the control, the multiple gRNA injected G0 animal and the G1 animal in Figure 4. Unmodified sequences include the reference sequence (first line) and others with small modifications (in red) away from the Cas9 predicted cleavage site. Modified sequences are those with substitutions or deletions (in red) at the predicted Cas9 cleavage site. The PAM is highlighted. gRNA target sequences in bold.

**Table S2.** Primers for *in vitro* gRNA synthesis. In bold the crRNA region aligning to genomic DNA.

**Table S3.** Sequencing primers used.

**Table S4.** Amplicon Seq analysis for *Rp*-*yellow* in y29 G0 nymph (Figure 2).

**Table S5.** Amplicon Seq analysis for *Rp-white* in G0 resulting from single *w1* gRNA injected female (Figure 3).

## Notes

### Competing Interest Statement

The authors have declared no competing interest.

